# Single-Cell Local Stress Analysis in Tumoroids

**DOI:** 10.1101/2024.01.22.576025

**Authors:** Rick Rodrigues de Mercado, Klara Beslmüller, Daan Vorselen, Erik H.J. Danen, Thomas Schmidt

## Abstract

The reciprocal interplay between cancer cells and their local environment, mediated by mechanical forces, necessitates a deeper experimental understanding. This requires precise quantitative measurements of cellular forces within the intricate three-dimensional context of the extracellular matrix. While methods such as traction-force microscopy and micropillar-array technology have effectively reported on cellular forces in two-dimensional cell culture, extending these techniques to three dimensions has proven exceedingly challenging. In the current study, we introduced a novel approach utilizing soft, elastic hydrogel microparticles, resembling the size of cells, to serve as specific and sensitive traction probes in three-dimensional cell culture of collagen-embedded tumoroids. Our methodology relies on high-resolution detection of microparticle deformations. These deformations are translated into spatially resolved traction fields, reaching a spatial resolution down to 1 µm and thereby detecting traction forces as low as 30 Pa. By integrating this high-resolution traction analysis with three-dimensional cell segmentation, we reconstructed the traction fields originating from individual cells. Our methodology enables us to explore the relationships between cellular characteristics, extracellular traction fields, and cellular responses. We observed that cellular stresses ranged from 10 to 100 Pa, integrating to cellular forces from 0.1 to 100 nN, which correlated with the localization of the cells actin skeleton, and the interaction area that cells developed towards the microparticles. Interestingly, the interaction of cells with inert microparticles appeared to be governed by contact mechanics resembling that of two soft spheres. The methodology presented here not only addresses the challenges of extending traditional stress-probe techniques to three dimensions, but also opens a strategy for the study of specific interactions between cells and the local tumoroid environment in a strive to further understand cell-matrix reciprocity in tissue.

Here, we present a novel methodology that permits the measurement of quantitative surface stresses on small, inert, elastic, deformable microparticles. Our approach tackles the involved task of mapping local three-dimensional stress fields within tissue. Our methodology was successfully applied to analyze local stresses within a tumor spheroid. We foresee that our research represents a significant advancement toward comprehending the intricate dynamics of cell-matrix reciprocity within tissue.

## Introduction

Metastasis is the primary cause of cancer mortality [1]. During cancer metastasis, cells dissociate from the primary tumor and spread to distant sites in the body. The metastatic potential of cancer cells often go hand in hand with therapy resistance [2]. All pathological signatures of metastasis, tumor growth, local tissue invasion, dissemination and therapy response are thereby controlled by biochemical as well as mechanical crosstalk between the tumor and its tissue microenvironment. An understanding of the mechanisms underlying such crosstalk in metastasis is hence crucial, and might lead to the potential development of novel lines of therapies. Here, we zoom in on the reciprocal mechanical interaction between tumor cells and their microenvironment that contributes to cell movement. Reciprocal interaction relates to the relation between active forces of cells, and the mechanical characteristics of the extracellular matrix (ECM) [3]. Essential questions to be addressed for understanding the mechanoreciprocity in cancer are, to what extent do cell-generated forces enhance their metastatic potential, in how far are the active mechanical properties of malignant cells different to those of non-malignant or healthy cells, and in how far are we able to modulate mechanical phenotypes of cells.

The role of mechanical properties of cancer cells [4] as well as that of the microenvironment, including cell or matrix stiffness, has only been studied in recent years. Most of the studies investigated the mechanical properties of tumor cells growing in cell cultures in a two-dimensional substrate dish. Methods, like traction-force microscopy either using flat soft gels [5], micropillar arrays [6], atomic-force microscopy [7, 8], and optical stretcher devices [9], are exquisitely able to report on the mechanical properties of individual cells and on cells growing in 2D. Extensions of these established methods to three dimensions rely on measuring deformations of the complex extracellular network surrounding the cells, the mechanical properties of which being highly non-linear and inhomogeneous, which makes the extraction of cellular forces in tissue extremely difficult [10]. Yet, metastasis formation indisputably is a process that occurs in the more complex environment of a three-dimensional tissue. It has been shown that force generation and migratory behavior of cancer cells in 2D differ significantly to those in which cancer cells are embedded in a 3D ECM [11, 12].

The ECM thereby plays an important role in promoting or inhibiting cell migration during cancer metastasis. The ECM is composed of a large ensemble of proteins, including fibrillar network-forming proteins like collagen or fibronectin. Tumors produce ECM components, proteolytically degrade ECM components, and bind and exert forces onto ECM components through adhesion receptors such as integrins [3, 13]. In turn, the chemical, mechanical, and structural features of the ECM fibrous network influence tumor growth and therapy response as well as tumour cell motility and invasion [14, 4, 15, 16]. For example, cancer cells often show increased collagen degradation and remodeling compared to normal cells, which in this way contribute to the formation of invasive tumors [17]. Conversely, it has been shown that the mechanical properties of collagen, such as the stiffness, may influence the migration of cancer cells.

In order to further understand the interaction between cells and their environment, 3D *in vitro* assays have been established in which tumor and ECM interact much alike to the situation in the corresponding tissue [18, 19]. These 3D culture systems closely mimic the *in vivo* pathophysiological situation in human tumor tissue. Spherical aggregates of tumour cells, so called multicellular tumor spheroids (tumoroids), are thereby embedded in reconstituted ECM environments [20]. Such reconstituted ECM environment give structural support for cell attachment and can be made from different types of elastic hydrogels [21], ideally with equivalent mechanical properties to the natural ECM.

In earlier studies we have established automated printing of cell clusters into 3D fibrillar ECM networks and used this approach to show how tumoroids exert forces onto the surrounding ECM to locally reshape their environment [20, 22]. Forces exerted by single cells embedded in 3D ECM have been studied by analyzing the displacement of small fiducial markers that were randomly embedded in the matrix before cross-linking the gel [23]. Such an approach had some success, yet needed a detailed knowledge of the local mechanical behavior of the fibrous network, being difficult to model. Therefore it is advantageous to have methodology at hand that permits determination of small cellular forces in three dimensions directly. One approach was the development of cell-sized “pressure sensors”, gel-beads with randomly embedded fiducial markers. [24, 25, 26, 27]. Displacement of the markers thereby report on the deformation of the sensor which was subsequently translated into local traction at the resolution of the pressure-sensor.

Here we present an advanced methodology that reports on local stress at the sub-cellular, micrometer, length scale inside of ECM-embedded tumoroid cultures. The methodology does not require prior knowledge of the mechanical characteristics of the fibrous environment, and still has a resolution that permits to assign local stresses to the action of individual cells. We utilized soft hydrogel microparticles, previously demonstrated to accurately capture the forces exerted by individual phagocytic cells on their substrate [28]. Here, we expanded upon this methodology to investigate the mechanical characteristics of cancer cells within collagen-embedded tumoroids. The microparticles, sized comparably to cells (20-50 µm), exhibit mechanically uniform and isotropic properties that are easily adjustable across a broad stiffness spectrum ranging from 100 Pa to 10 kPa. This versatility covers the typical stiffness range of various cells and tissues. Their size enables precise stress analysis at the micrometer scale. By combining local stress measurements on these microparticles with a sophisticated cell-segmentation algorithm, we were able to accurately attribute localized stresses within the three-dimensional matrix of a gel-embedded tumoroid to individual cells. Our innovative approach facilitates the investigation of directional forces within the intricate three-dimensional context at the single-cell level, enabling the measurement of cellular forces transmitted through the collagen matrix.

In our study we aimed to characterize the interactions of individual cells with their neighbouring cells inside a tumoroid, and of individual cells with the collagen fibers of the ECM network. Microparticles were coated with bovine-serum albumin which prevented specific interactions of cells with the particles, and rendering the particles as passive probes for local stresses. Forces generated by cells resulted in a local stress-field as observed by local deformations of the microparticles. The homogeneous mechanical properties of the microparticles, subsequently allowed us to quantitatively translate the microparticle deformations into traction-fields at the particles surface to a resolution of ≈1 µm and traction fields as low as ≈30 Pa. Our analysis was combined with a 3D cell-segmentation. The accurate three-dimensional shape of individual cells in an image stack was determined. Together with the segmentation we obtained a detailed, cell-centered traction-field, which permits us to associate extracellular traction fields, and cellular response to a quantitative phenomological description of cell behavior.

## Materials and methods

### Microparticle synthesis

Microparticle synthesis and functionalization was performed similar to previously published [28]. Deformable acrylamide-co-acrylic-acid microparticles were synthesized using Shirasu Porous Glass membranes (SPG Thechnology, Japan). These tubular membranes with tunable pore size enabled us to produce spherical particles in large quantities and of uniform size in a range of 5-50 µm. Before use, the SPG-membrane was submerged in HPLC-grade N-heptane (Merck, 1043792511) and placed in a vacuum chamber to remove air bubbles. After sonicating the membrane for 5 min in N-heptane, the membrane was mounted into the internal pressure micro kit extruder (SPG Technology). A solution of 200 mL hexane (Fisher Scientific, Hampton, NH, USA, P216885) with 3 % (w/v) surfactant span80 (Fluka, 85548) was prepared and poured into a 3-neck water-jacketed flask together with a magnetic stirrer bar. This mixture defines the continuous phase. The flask was connected to a water bath, a temperature sensor was inserted, put under a nitrogen sparge, and placed on a hotplate. The solution was stirred at 350 RPM.

A solution of 150 mM NaOH, 0.3 % (v/v) tetramethylethelenediamine (TEMED; Thermo Fisher, Waltham, MA, USA, 17919) and 150 mM MOPS sodium salt (Sigma-Aldrich, Waltham, MA, USA, M9381) was supplemented with acrylamide (AAm), acrylic acid (AAc) and cross-linker N,N’-methylene-bisacrylamide (BIS) (all Sigma-Aldrich, A9099, 147230, 146072, respectively) with a final pH of 7.4. Two acrylamide mixtures were used for particle synthesis. The total mass concentration of acrylic components *c*_*tot*_ = *c*_*AAm*_ + *c*_*AAc*_ + *c*_*BIS*_ = 100 mg/mL for both mixtures. The relative concentration of acrylic acid was set to 10 % for the first mixture. The second mixture had a relative concentration of acrylamide of 10 %. A cross-linker concentration *c*_*c*_ = *m*_*BIS*_*/*(*m*_*AAm*_ + *m*_*AAc*_ + *m*_*BIS*_) of 0.65 % and 1.5 % was used for the first and second gel mixture, respectively. The Young’s modulus of the particles was characterized using atomic force microscopy. The first set of particles had a Young’s modulus of *E* ≈ 1 kPa, the second set of *E* ≈ 0.6 kPa.

The acrylamide solution (dispersed phase) was degassed for 15 min. 10 mL of the dispersed phase was transferred to the reservoir of the SPG module. The reservoir was subsequently flushed with nitrogen. The critical pressure for particle generation was estimated from the difference in surface tensions of the two liquid-mixtures [28]. The starting pressure was set just below this value. The pressure was then slowly increased, until a flow was observed. The acrylamide mixture was allowed to be pushed through the membrane for 3 h. Subsequently, the SPG module was retracted, the flask was plugged and the temperature increased to 60 °C. Polymerization was initiated by adding 1.5 mg/mL 2,2’-Azobisisobutyronitrile (AIBN; Sigma-Aldrich, 441090) to the solution. After 3 h at 60 °C, the temperature was reduced to 40 °C. After 16 h, the particles were finally collected in round-bottom tubes and washed thrice in hexane, followed by a wash in ethanol. All ethanol was evaporated and particles were stored in PBS containing 5 mM sodium azide (Sigma-Aldrich, S20002).

### Microparticle functionalization

First, particles were washed in activation buffer (100 mM MES (Sigma-Aldrich, M5057) and 200 mM NaCl), and subsequently incubated for 15 min in reaction buffer of 0.1 % tween20 (Sigma-Aldrich, P7949), 4 % (w/v) EDC (Sigma-Aldrich, E7750) and 2 % (w/v) NHS (Thermo Fisher, 22500). Then, the microparticles were washed and incubated for 1 h with 5 µg/mL of BSA in PBS at pH 8. To verify the efficiency of protein binding, Alexafluor-488 conjugated BSA was used. Fluorescent images confirmed the protein binding (Fig. 1 A). Subsequently, TexasRed-Cadaverine (Thermo Fisher, T2425) was added for 30 min. Unreacted NHS groups were blocked using 300 mM Tris and 100 mM ethanolamine (Sigma-Aldrich, 398136) (pH 9). Finally, particles were washed thrice in PBS with 0.1 % tween20 and stored for use in PBS (pH 7.4) with 5 mM sodium azide.

**Figure 1:**
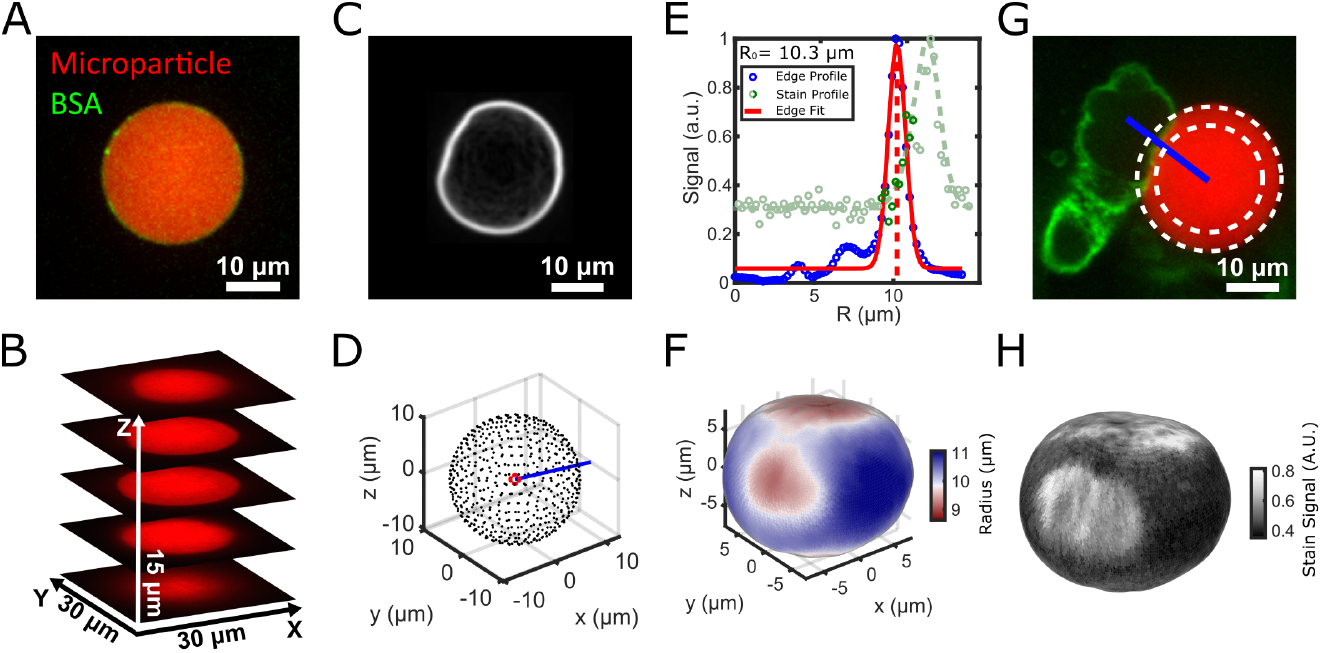
Particle shape and signal reconstruction. **A**. BSA conjugated with AlexaFluor-488 confirmed adhesion of BSA to the particle’s surface. The particle was stained with TexasRed-Cadaverine. **B**. A z-stack of the particle was taken. **C**. Edges were found using a Sobel operator. **D**. Lines were drawn from the center of the particle radially outward. **E**. A Gaussian fit (red) was used to find the peak of the edge line-profile (blue). The same lines were used to find the line-profile of the stain signal (green). The stain signal on the particle’s surface was calculated by averaging the signal within 1 µm radius of the edge (dark green). **F**. Triangulation was used to reconstruct the shape. **G**. The stain (AlexaFluor-488 Phalloidin, green) intensity of MV3 cells was found by drawing lines (blue) from the center of the particle (red). **H**. The signal on the particles’ surface was averaged and projected onto the sphere-like shape.

### Atomic-force microscopy

The Young’s modulus of the particles was determined by atomic-force-spectroscopy on a JPK NanoWizard 4 atomic-force microscope (AFM; Bruker, Berlin). A plastic dish was activated using UV-ozone for 10 min. Particles were incubated for 15 min in activation buffer (see section “Microparticle functionalization”) with 0.1 % tween20, 4 % (w/v) EDC and 2 % (w/v) NHS. Unbound particles were removed by washing with PBS. Measurements were performed in PBS at about 25 °C. For the measurements, spherical-shaped probes (Nanoandmore GmbH, Germany) were used (CP-qp-CONT-BSG-B-5) with a nominal spring constant of 0.1 N/m, and nominal tip diameter of 10 µm. The cantilevers were calibrated before each measurement with the thermal tuning method [29, 30], and also inspected with scanning electron-microscopy (SEM) for accurate determination of cantilever dimensions and sphere radii. For determination of the elastic modulus we employed the shallow indentation method. Using an optical microscope, the spherical probe was centered on the particles. The indentation curves where performed to a force-maximum of 1 nN. To reduce noise, 4-5 nano-indentations were performed on each bead. Force-distance curves were corrected for cantilever bending. Each FD curve was transformed into a force-separation (FS) curve (Fig. S1). This curve reflects only the sample indentation, excluding thereby the effect of the cantilever bending. The Young’s modulus was obtained by fitting the FS-curves to the Hertz model with spherical indenter [31].

### Cell culture

The human melanoma cell line MV3 [32] was cultured in high-glucose Dulbecco’s Modified Eagle’s Medium (DMEM, Gibco, 11504496) containing L-Glutamine and Sodium Pyruvate, supplemented with 10% fetal calf serum (Thermo Scientific) and 100 µg/mL penicillin/streptomycin in a humidified incubator at 37 °C with 5% CO_2_.

### 3D collagen matrix

1 mg/mL collagen gels were created by mixing rat tail collagen type I (5 mg/mL stock, Biotechne, 3447-020-01) with buffers. Two different buffer types were used and compared: for HEPES buffering, we mixed DMEM (Gibco, ThermoFisher Scientific, Waltham, MA, USA), HEPES 0.1 M (1M stock, Biosolve, Valkenswaard, The Netherlands) and NaHCO3 44 mM (440 mM stock, Merck, Darmstadt, Germany) as previously described [33]. For phosphate buffering we mixed disodium hydrogen phosphate (Sigma Aldrich, Cat. No. 71636) and sodium dihydrogen phosphate (Sigma Aldrich, Cat. No. 71507) to pH value 7.4 and final phosphate concentration of 200 mM as described before [34]. Fluorescent particles were added to the collagen mixture to a concentration of 80p/uL. For the HEPES buffered collagen gel, fluorescent particles with a Young’s modulus of 1 kPa were used, and for phosphate buffered collagen gel fluorescent particles of 0.6 kPa were used. The collagen-particle mixture was polymerized in a 96-well Sensoplate (Greiner Bio-one, 655892) at 37 °C for 1 h.

### Tumoroid formation

Tumoroid formation into 3D collagen matrix was performed as described previously [20]. Cells were mixed with fluorescent microparticles in a ratio of approximately 1000:1 and resuspended in 2 % polyvinylpyrrolidone (Sigma-Aldrich, P5288) to reach a cell concentration of 25000 cells per µL shortly before cells were injected into collagen. Tumoroids, 200 µm in diameter, were created by automated injection of cell-particle mixture into the collagen gels using the injection robotics from Life Science Methods (Leiden, The Netherlands). After injection, tumoroids were incubated with the appropriate medium at 37 °C. After 24 h, tumoroids started to invade into the 3D collagen matrix. Samples were subsequently incubated in a fix-stain solution containing final concentrations of 2 % formaldehyde (Sigma-Aldrich, 252549), 0.1 % TritonX-100 (Sigma-Aldrich, T8787), 0.4 µg/mL Hoechst (Fisher Biotech, 33258) and 0.05 µM AlexaFluor-488 Phalloidin (Invitrogen, A12379) in PBS for 3 h at room temperature. The samples were washed thoroughly in PBS and measurements were performed shortly after.

### Microscopy

#### Scanning confocal microscopy

Images of tumoroids were acquired on a Nikon Eclipse Ti inverted scanning confocal microscope equipped with four laser lines 405 nm, 488 nm, 561 nm and 640 nm, and with an A1R MP scanner. The microscope has a Nikon encoded and automated stage. Its camera is controlled through NIS Element Software (Nikon Instruments Inc., Melville, NY, USA). Tumoroids were imaged with 20 µm distance between Z-slices. ECM collagen-fibers were detected by confocal reflection microscopy on the same scanning confocal microscope. The collagen fibers were scanned at 561 nm excitation with a 561 nm blocking dichroic mirror and all the light reflected back passed a through filter of bandwidth 400 nm - 750 nm. The reflected light from the collagen fiber network was collected on GaAsP-photomultiplier. Scanning confocal fluorescent microscopy of tumoroids, and scanning confocal reflection microscopy of the collagen network were performed using Plan Apo ×20/0.75 NA, and Apo LWD ×40/1.15 objectives (Nikon Instruments Inc., Melville, NY, USA), respectively.

#### Spinning-disc confocal microscopy

High-resolution imaging was performed on an inverted microscope (Zeiss Axiovert 200) equipped with a 63X, 1.15 NA water LD C-Apochromat W korr objective (Zeiss) of 600 µm working distance. The setup was expanded with a confocal spinning-disk unit (Yokogawa CSU-X1), and an emCCD camera (Andor iXon DU897). The hydrogel microparticles were imaged with 561 nm DPSS-laser illumination (Cobolt). A 488 nm DPSL-laser (Coherent) was used to image AlexaFluor-488 Phalloidin. A 405 nm diode laser (CrystaLaser) was used to illuminate the Hoechst nuclear-stain. Z-stacks were used to obtain a 3D image-cube of the microparticles and the surrounding cells. The mismatch between the refractive indices of the immersion-oil of the objective and the culture medium caused an apparent elongation of the image cube, which was corrected by experimentally-determined correction factor: the apparent elongation of more than 10 stiff (*>* 5 kPa) microparticles was determined by calculating the radius in the equatorial plane compared to the lateral radius of the particles. The ratio of both was used as a correction factor for the z-positions, rendering the effect of the refractive index mismatch negligible.

### Local traction analysis

Fast traction analysis was performed in spherical harmonics space [35] implemented in Python’s SHTools library [36]. Spherical harmonics are solutions to the generalized linear elasticity continuity equation (Hooke’s law) for the mechanical equilibrium condition of a spherical boundary in an isotropic medium. Hooke’s law in terms of the displacement field **u** reads:

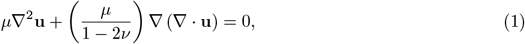

where *µ* is the shear modulus of the spherical object (microparticle), and *ν* is the Poisson’s ratio of the elastic medium. The displacement field in turn is expressed in complex coefficients, 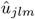, of the spherical harmonics:

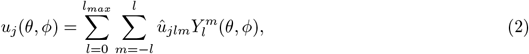

where 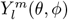 are the set of complex spherical harmonics functions given by the harmonic-degree *l* and order *m*. The displacement field results in a traction field, *T*_*j*_(*θ, ϕ*):

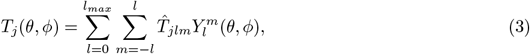

where the components 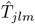 are uniquely defined by 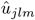.

The proper displacement field, 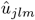, that solves Eq. (1) is found iteratively by minimizing the total elastic energy, and the total traction on the particle for which the deformations pointed outward. The latter choice rooted in the fact that cells were unable to pull on the surface-passivized particles. It should be noted that omitting this latter requirement solely introduced an offset in the traction without changing the overall shape of the traction-field on the particles (Fig. S2).

Any local stress on the microparticles led to a local deformation of the particle’s surface at the location of the stress, and to a small global deformation of the particle due to the counterbalancing action of the embedding ECM. In some cases this counterbalancing action was directly observed as local deformation along a thick ECM-fiber close to the particle’s surface (Supplemental Fig. S3).

The spatial resolution of the traction field is limited by the maximum harmonic degree, *l*_*max*_, in the spherical harmonics expansion. *l*_*max*_ was chosen based on the trade-off between computational power and desired spatial resolution. The spatial resolution, Δ*r/R*, scales almost linearly with *l*_*max*_ (see supplemental Fig. S4)

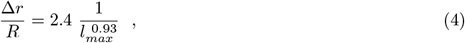

where *R* is the radius of the spherical object. In our study we used *l*_*max*_ = 25, since this gives a good balance between spatial resolution (≈1 µm), and computational time (≈5 h per particle on an office PC).

### Statistics

The Mann-Whitney U-test was used to determine statistical significance between populations. Data sets were significantly different with probabilities of *p <* 0.001 (***), and not significantly different for probabilities of *p >* 0.05 (ns).

## Results

### High-resolution microparticle shape reconstruction

The equilibrium shape of the elastic microparticles is that of a sphere. Application of a force to the particle will result in its deformation. Hence, the higher the resolution that the particle’s surface is reconstructed, the smaller the minimum forces are that we are able to detect. Here we developed a high-resolution 3D particle-surface reconstruction methodology that allowed us to reconstruct the particle shape to the statistical limit set by the signal-to-noise ratio of detection.

All particles were homogeneously labeled by fluorophores during the preparation step of the particles (see M&M). Z-stacks of particles were taken on a spinning-disk confocal fluorescence microscope. The z-slices were separated by at a distance of Δ*z* = 370 nm which is half the depth-of-focus of the objective, close enough to comply to Nyquist’s theorem. Care has been taken to adjust the excitation intensity and illumination time such, to keep the emCCD-detector out of saturation. Typically image-cubes of 112 *×*112*×* 80 µm^3^ were acquired. Individual particles in the image-cube were subsequently identified, and the data cubes cropped to contain solely individual particles (Fig. 1 A,B). These sub-cubes were further analyzed following an analysis pipeline similar to that described by Vorselen *et al*. [28].

In short, the particle’s edge was highlighted by applying a 3D Sobel-operator to the image cube (Fig. 1 C). Subsequently, radial line profiles of the filtered edge-data and radiating from the particles’ center-of-mass were calculated using cubic interpolation (Fig. 1 D,E). Line profiles were calculated on a regular grid of a perfect sphere at a distance equivalent to half the point-spread function of the microscope. Each of the radial profiles were fit to a Gaussian, returning the position of the edge, its width, and its integrated signal (Fig. 1 E). The local radius of the particle at the position shown in the line profile was 10.3 µm. The width returned from the edge-fit was, as predicted, equal to twice the width of the point-spread-function of the microscope (≈350 *nm*). The high-resolution edge positions, all spaced on a regular grid on a sphere, were ultimately used for the high-resolution reconstruction of the particles’ shape using triangulation (Fig. 1 F). The particle shown in (Fig. 1) had a mean radius of *R*_0_ = 10.1 µm. Determination of the high-resolution particle shape was exquisite. Analysis of ideal spherical particles on a coverslip and embedded in a buffer solution showed a root-mean-squared surface roughness of 50 nm, equivalent to the predicted edge localization error set by the photoncount level, and the width of the point-spread function (Fig. S5, S6) [37].

In Fig. 1 F we clearly identified particle deformations of up to 1 µm, way beyond the optical resolution and the potential surface roughness. The particle shown in the figure had a nominal radius of 10.1 µm. The nominal radius was determined from the particle’s total volume, as we assumed that the microparticle was incompressible and that volume was conserved when the sphere was deformed. The colors in Fig. 1 F indicate the sign of the deformation: red indicates indentations, blue protrusions.

Simultaneous with the edge detection, cells in close neighborhood to the microparticle were detected. Cellular signals were identified through labeling cells with an fluorescence stain of actin (Fig. 1 G). The actin signal (green in Fig. 1 E) radially located within 1 µm of the particles edge was summed (Fig. 1 E, dark green data points) to obtain an indicator for cellular vicinity, and actin concentration. The resulting stain signal-profile was then projected onto the edge-coordinate of the particle (Fig. 1 H). Clearly visible in Fig. 1 is that regions of high stain signal (Fig. 1 H) were colocalized with the indentations of the particle (Fig. 1 F). Cells obviously pushed on the particles which lead to the local particle deformations observed.

The goodness of the microparticle shape reconstruction depends on the local signal to noise ratio (SNR). Poor shape reconstruction originates from low SNR and erroneous Gaussian fitting of the microparticle’s edge as a consequence. The resulting spherical harmonics displacement field is unable to capture the extreme fluctuations of the erroneous fits. Hence to correct for these errors, a global goodness-of-fit estimator, *χ*^2^, for each particle surface reconstruction was calculated:

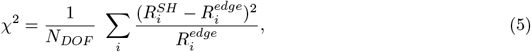

where, *N*_*DOF*_ = *d* · (*l*_*max*_ + 1)^2^, are the number of degrees of freedom in the fitting routine, with *d* the number of dimensions, 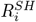, the radius at position *i* of the spherical harmonics displacement field, and, 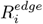, the radial edge coordinate from the Gaussian fitting routine. Only particles of *χ*^2^ *<* 0.5 were considered for further analysis. By this criterion *<* 10% of the particles were removed from the dataset.

### Microparticles in ECM-embedded tumoroids

Fluorescent microparticles were used in tumoroid assemblies, in which tumoroids together with particles were embedded within a collagen matrix, resembling a native ECM environment. The ECM surrounding tumoroids consists of a network of collagen fibers that each were approximately 50-100 µm in length (Fig. 2 C). The assembly was established by microinjection of a cell / microparticle suspension into a preformed gel. 24 h after injection MV3 cells had migrated into the ECM and away from the injection site (Fig. 2 A). Z-stacks of tumoroids were taken on a confocal microscope. Multiple particles were located within the ECM-embedded tumoroid. The microparticles were coated with bovine serum albumine, which rendered them inert with respect to cell interactions. The particles localized at random into the tumoroid (Fig. 2 B).

**Figure 2:**
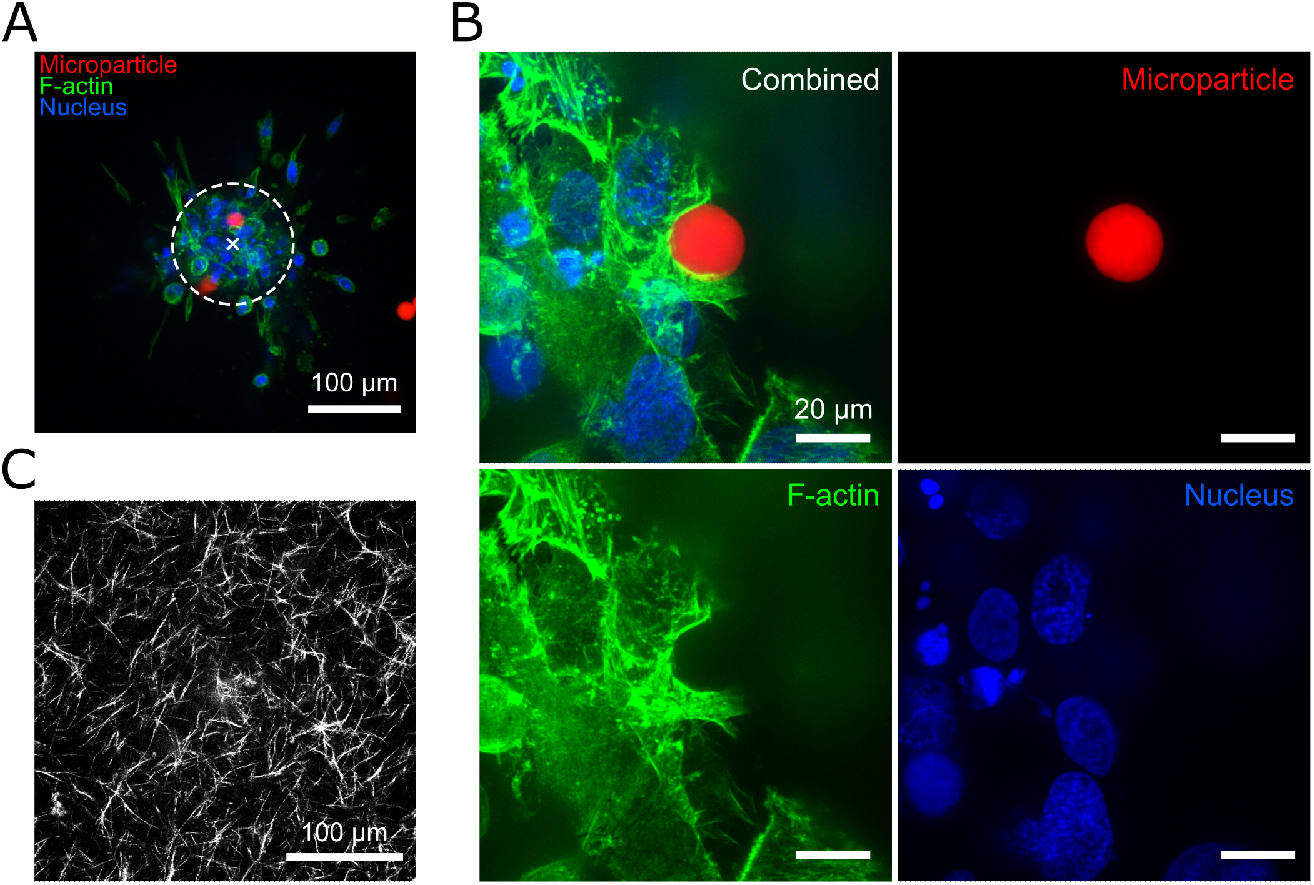
Tumoroid with particles in collagen gel, 24 h after tumoroid injection. **A**. MV3 tumoroid with cells migrating from the injection site (marked with ‘x’). The dashed line indicates the tumoroid core. F-actin was labelled with AlexaFluor-488 Phalloidin and nuclei with Hoechst. **B**. Close-up of a particle at the edge of a tumoroid. All scale bars are 20 µm. **C**. Reflection microscopy image of the collagen network.

### Single-cell traction decomposition

In Fig. 1 F-H it appeared that indentations were most prominent where cells were localized close to the particle. For quantitation of this observation we developed methodology that automatically identified cells, and allowed us to correlate particle deformations with various cellular properties. Our methodology identified regions on the particles surface that were in proximity to individual cells, from which we deduced from each cell the amount of the force it applied to the particle.

#### Cell segmentation

Cell segmentation inside a tumoroid within the 3D image-cube was developed based on an analysis pipeline described by Grosser *et al*. [38] (supplemental Fig. S7). The method required both the nucleus and the cell-membrane to be stained. In brief, an adaptive threshold was used to generate a 3D-mask of all nuclei in the image cube. The mask was dilated, holes were filled and subsequently eroded. Neighbouring nuclei were separated using a watershed algorithm. The nuclear centers were dilated to a size smaller than a typical nucleus (≈2 µm), serving as seed in the thresholded channel of the cytoskeletal actin defining the cell’s cytosol, and resulting in the final segmentation of the image cube. Potential lower fluorescence levels at the cell edges were cut out by subtracting a morphologically opened image. Finally, salt-and-pepper noise was reduced by applying an adaptive median filter.

In addition, the boundary of the tumoroid was identified by thresholding a blurred, entropy-filtered image of the cell-edges. A Sobel filter was used to find the the cell edges, which were subsequently smoothed with a narrow kernel to merge peaks in the gradients of the same line segment. Finally, the image was superimposed with the nuclear seeds and watershed segmentation was performed. Such watershed segments are displayed in white in Fig. 3 A.

**Figure 3:**
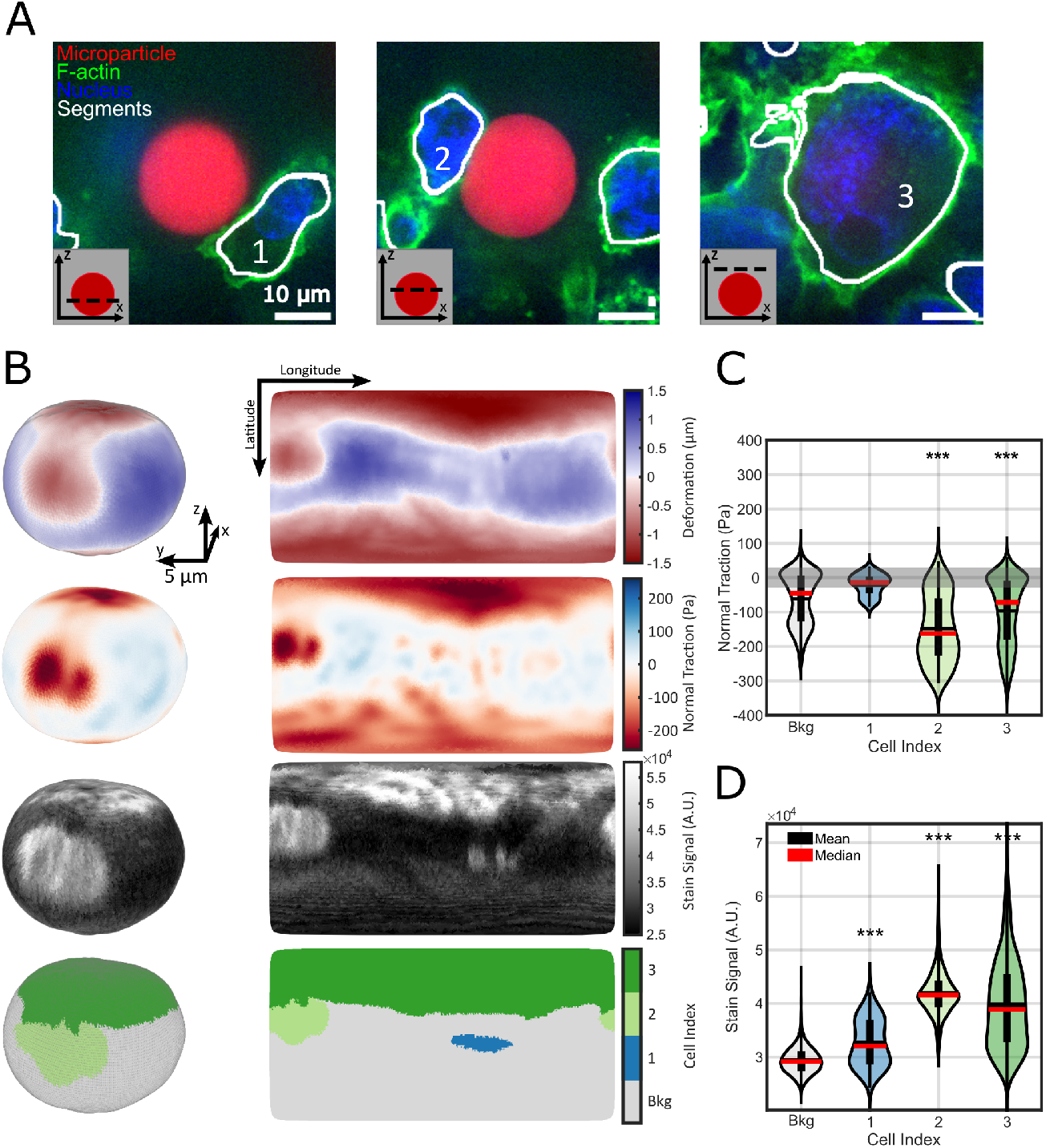
Cell segmentation in tumoroids allows for single-cell traction-analysis. **A**. Slices of a z-stack of a MV3 tumoroid in a collagen gel. The particle (red) was located at the edge of the tumoroid. The cell shape was obtained by staining F-actin (green). Cells were identified by their nuclei (blue). Here, three cells were found touching the particle. The obtained cell-segments are displayed in white. **B**. From top to bottom: the deformation, normal traction, stain signal and cell index map. The cell indices correspond to the labels in (A). Unexposed regions of the particle were identified as background (bkg). **C**. Cells 2 and 3 significantly indent the particle, hence applying a negative normal traction beyond our traction resolution (grey area) (*p <* 0.001). **D**. The particle surface that was exposed to cells had an increased amount of stain signal (*p <* 0.001).

The quality of the fluorescent signal of both the nuclei and cell membranes determine the segmentation accuracy. Simulated data at equivalent signal-to-noise values to our measured image stacks showed, that *>* 80 % of the line segments were correctly detected.

#### Single-cell traction analysis

In Fig. 3 A three melanoma MV3 cells (indexed 1-3), each at different z-position with respect to a central microparticle, were identified to interact with the single microparticle shown. The microparticle was localized at the border of a collagen gel-embedded tumoroid. To characterize the force each of the cells applied on the microparticle, their nuclei (blue) and cytosol (actin signal, green) were segmented as described above. The cell-segments are indicated in white in Fig. 3 A.

From the particle deformations (Fig. 3 B top) the traction field normal to the surface of the particle (Fig. 3 B 2^*nd*^) was determined as further described in the M&M section. The normal traction ranged from -200 Pa (inward, red) to +50 Pa (outward, blue). Further fig. 3 B shows the locations of the nearby actin-skeleton of cells (3^*rd*^ row), and the respective cell indicators (1-3, bottom). In the bottom panel, regions on the particle unexposed to cells were identified as background (indicator “Bkg”, grey).

As already noted in Fig. 1, indentations of the microparticles were visible where cells touched. Cells obviously applied pushing forces (negative normal traction) to the soft microparticles. As the particle shown was located near the periphery of the tumoroid, the largest part of the surface (861 µm^2^, 66%) was not covered by cells. The contact areas of cells varied between 27 µm^2^ (2%, blue cell “1”) to 66 µm^2^ (5%, light green cell “2”), and 348 µm^2^ (26%, dark green cell “3”). At areas where cells were in proximity to the particle, the traction normal to the surface was significantly different to those areas not covered by cells (“Bkg”) (*p <* 0.001). The mean of the absolute normal traction varied between 29 Pa (blue, cell “1”), to 143 Pa (light green, cell “2”), and 71 Pa (dark green, cell “3”) (Fig. 3 C). By combining cell segmentation with traction analysis our method thus allowed us to quantitatively describe the mechanical interactions that cells within tumoroids develop towards their direct tissue-like microenvironment.

Combining the values for the mean cell traction with those we obtained for the respective cell area touching the microparticle, we calculated the total force that each cell exerted on the particle. Those forces varied between 0.60 nN (blue, cell “1”), to 9.0 nN (light green, cell “2”), and 21 nN (dark green, cell “3”), respectively. It should be noted, that clearly overall force-balance holds. The microparticles were about 10-fold bigger than the pores in the collagen gel, hence force balance is faithfully guaranteed by a counter force of the gel being spread over the full surface of the particle. These counterbalancing forces are the reason for the exceeding spread of positive traction in the background region in Fig. 3 C (grey), as compared to the control measurements of microparticles in buffer solution.

We further analyzed the F-actin signal for each of the cells as a means to quantify the potential activity of the cells. The mean actin signal of cell “1” was significantly smaller than that of the two other cells touching the particle (Fig. 3 D). By comparing Fig. 3 C with Fig. 3 D it appeared that the mean traction correlated with the amount of actin in the contact area of cells with the particle. Such a correlation is to be predicted, since F-actin is the main force generating machinery cells have to develop active forces on their environment.

### Cellular forces increase with F-actin density and interaction area

We wanted to verify the correlations we observed for the individual particle shown in Fig. 3, and test the hypotheses that cells individually develop forces inside the tumoroid. The predictions include that cells apply larger forces when they have a more pronounced actin skeleton, and that cells apply more forces when they are able to make more contact with their environment. Both dependencies have been observed for cells grown on a 2D substrate [39, 40, 41, 42, 43, 44], yet are still to be shown for the complex environment of a tumoroid.

For quantification of the level and density of the F-actin network, the F-actin signal, *S*_*act*_ needed to be quantified. Yet, the F-actin signal and the unavoidable background signal of each cell varied considerably depending on the location of the particle relative to the core of the tumoroid. Such variation is inherent to the measurements, and relies among others on the local labeling efficiency, the local intensity of the excitation light (shadowing), and on the local efficiency of the fluorescence detection. These parameters not only vary depending on the location within the tumoroid, but also from sample to sample. Yet all the above mentioned unknowns influence the signal inside the actin-free microparticle, *S*_*bkg*_, in an identical manner. Thus, we here define the reduced actin signal *S*^*^:

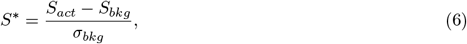

which corrects for experimental uncertainties and faithfully reports on the F-actin signal for all particles. The standard deviation of the background signal *σ*_*bkg*_ is thereby taken as reference. This rescaling to *S*^*^ equalizes the background for all images, and allows for quantitative comparison of the actin signal in different experiments.

In total, 55 + 43 cells interacting with 17 + 9 microparticles in 2 independent experimental runs were used for further analysis. The ensemble data verified the correlations observed for the individual particle presented in Fig. 3. Cells expressing more F-actin (increase in *S*^*^), applied more pushing on the particle (decrease in normal traction, Fig. 4 A). Cells for which the actin signal was similar to that of the background (*S*^*^ ≈ 0) did not develop local traction. In summary, traction significantly increased above background for cells in which F-actin was prominently detected.

**Figure 4:**
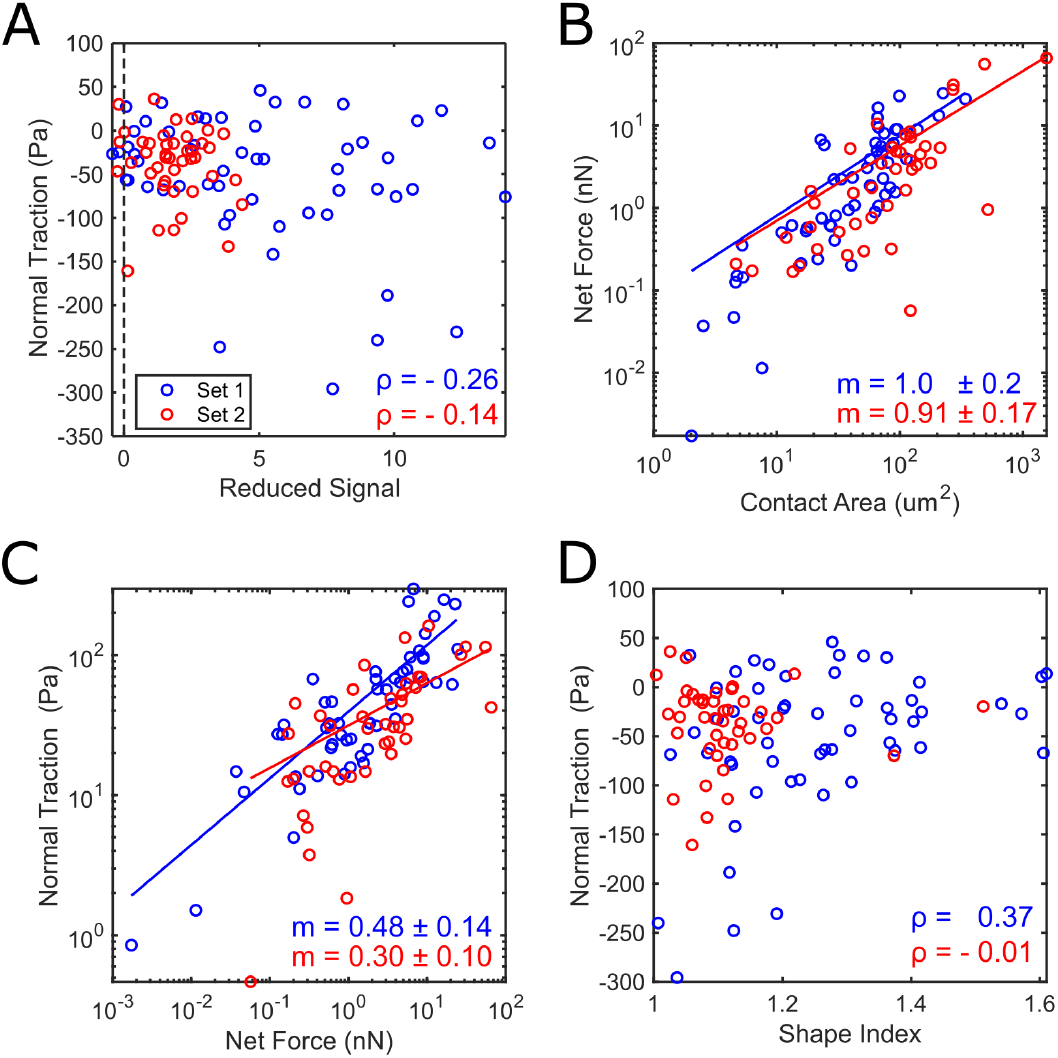
Contact area and the amount of actin correlate with the force cells apply. **A**. Weak correlation between mean normal traction at the cell-particle contact area, and reduced F-actin signal, for two independent experimental conditions (blue and red). Correlation coefficents, *ρ*, are indicated. **B**. The net force and contact area are strongly correlated through a power law relation. **C**. A power law relationship of slope *m* ≈ 1 was found between normal traction and net force. **D**. The applied normal traction appeared to be independent of the cell shape. Correlation coefficients are displayed by *ρ*. Power law fits are displayed by solid lines in (B-C). The power law exponent of *y* = *kx*^*m*^ including the standard error of the fit is displayed in the corner.

Further, we investigated how the forces that an individual cell exerted varied with the interaction area the cell had with the particle. The net force an individual cell applied on a microparticle was calculated by integration of the normal traction over the interaction area. Results are shown in Fig. 4 B. With increasing interaction area the magnitude of the net force increased. The dependence appeared to follow a power-law as judged from the linear scaling in a double-logarithmic representation. From the slope, *m*, in double-logarithmic representation (*m* = 1.0, cq *m* = 0.91) it appeared that the net force scaled proportional (*m* = 1) to the interaction area. A power-law relation was likewise observed between the average traction and the net force (Fig. 4 C). From the fit to a linear dependence in the double-logarithmic representation (*m* = 0.48, cq *m* = 0.30), we found that the normal traction increased with about the square root of the force. Given the experimental uncertainties in those experiments, both observation comply with a model that describes the contact mechanics of two elastic spheres, as further detailed in the discussion section.

Cell-segmentation allowed us further to determine the cell morphology and cellular orientation. Given that morphology and orientation of cells was shown to predict cellular motility [45, 46, 47, 48], we hypothesized that cells which were oriented head-on to the particle’s surface would develop a stronger normal traction on the particle, as compared to cells passing by, and likewise that cells passing by would develop an increased shear-traction on the bead. Overall cell morphology is typically expressed in terms of the cell shape-index, *s*, which is the ratio of the cell’s radius determined from it’s volume, *r*_*V*_ = 3*/*4*π V* ^1*/*3^, to that determined from it’s surface, 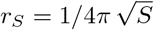[38]

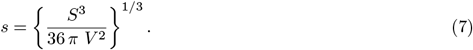

A perfect spherical cell would have *s* = 1, larger values indicated more elongated cells. Here we take the cell shape-index as an indicator of the elongation of a cell. Fig. 4 D shows the relation of normal traction as it varies with cell shape-index. Unlike our prediction, we did not find a correlation between shape index and normal traction (Fig. 4 D). Neither we find a correlation with the tangential traction of cells (see supplemental Fig. S8).

## Discussion

In the current study, we present a novel approach for quantitative analysis of local cellular forces in complex three-dimensional tissue-mimicking environments. Our method entails the combination of bulk synthesis of elastic microparticles, with high-resolution microscopy, and traction analysis inside a three-dimensional *in vitro* tumoroid experiment. By incorporating a cell segmentation pipeline into our analysis, we were able to address questions regarding cell-based inhomogeneities in stress generation inside of a tumoroid.

Cells developed stresses between 0.1 and 100 Pa and local forces between 0.1 and 100 nN, both compatible with the compliance of the gels used (≈500 Pa), and typical for values reported for various tissue [49, 50, 23, 10]. We predicted that cells apply larger forces when they produced more actin, and when they were able to make larger contact with their environment. We were unable to draw definitive conclusions regarding this prediction based on the observed results. While there are indications of a potential relationship between F-actin expression levels and measured stress, the data did not provide sufficient evidence to confirm the hypothesis.

The net force exerted by cells on the microparticles increased with increasing interaction area, where the net force scaled about linear with the interaction area. These findings are not too surprising when compared to the well-known problem of two soft spherical bodies in mechanical contact [31, 51]. From Hertzian contact mechanics, a similar trend is predicted in which the force *F* of elastic spheres of average radius *R* and effective modulus *E* increases with the final contact area *A* as

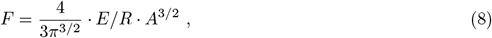

hence scales with the contact area to the power 1.5 [31, 51]. Likewise our finding that the normal traction scales with the square root of the net force come close to that predicted by the Hertzian model which reads that the pressure at contact *P*_0_, which we take as equivalent to the normal traction in our experiments, is given by,

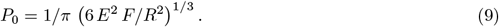

The deviations of our results from those ideal mechanical model predictions might be due to the crude assumption of a spherical cell for which the Hertzian models hold. It is worth to mention as conclusion that the magnitude of forces, in the nN-range, exerted by individual cells in the complex environment of a 3D matrix are consistent with those reported for cells plated on 2D surfaces obtained in traction force microscopy [52, 39, 53, 54, 44] and micropillar array techniques [43].

Surprisingly though, our study did not reveal a relation between cellular traction and cell morphology. Our data suggest that traction is independent of cell shape and orientation, in contrary to what we predicted from experiments we performed for cells in two-dimensions in a culture dish [48]. This latter result indicates that in complex three-dimensions environments further factors will determine cellular forces. Those potentially include specific cell-ECM interaction sites which could be realized by covalently attaching ECM components to the microparticles in future experiments.

Overall, our results suggest that cells within a tumoroid individually push on their environment, and that the degree of contact with other cells in their environment as well as expression level of cytoskeleton proteins affect the magnitude of the force exerted by the cells. Our study describes a new approach to characterize cellular forces in the context of their complex three-dimensional environment of the ECM. Here we provided first insights into the mechanics of cellular forces and and how those relate to tumor tissue mimicry. Our findings may represent a solid steppingstone for the use of the presented methodology in our strive to contribute to a better understanding of the cellular mechanics involved in tumor growth and metastasis.

## Acknowledgments

We thank Aditya Aniruddha Sane for assistance with setting up the microparticle fabrication.

## Author Contributions

R.R.M., K.B., E.D. and T.S. designed the experiments. R.R.M. and K.B. conducted the experiments. R.R.M. performed data analysis. R.R.M., K.B., D.V., E.D. and T.S. wrote the manuscript. All authors reviewed and edited the manuscript.

## Supplemental Material

**Figure S1:**
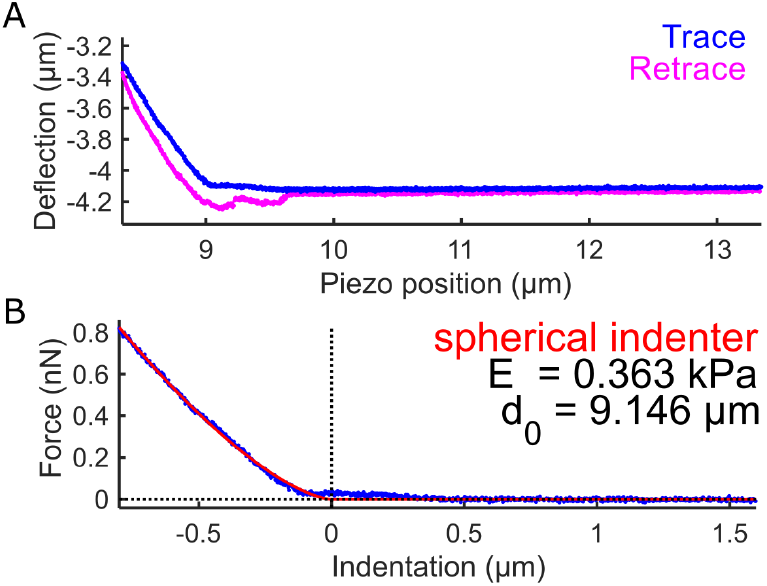
AFM force spectroscopy shows elastic Herzian contact mechanics. **A**. Unprocessed AFM force-indentation curve of a trace (blue) and retrace (magenta) scan from a spherical indentation probe on one of the microparticles. The indentation experiment was performed in buffer solution. **B**. Processed curve (blue) shows an indention up to 0.8 µm. A fit of the Herzian model is shown in red. This microparticle had a Young’s modulus of 0.36 Pa.

**Figure S2:**
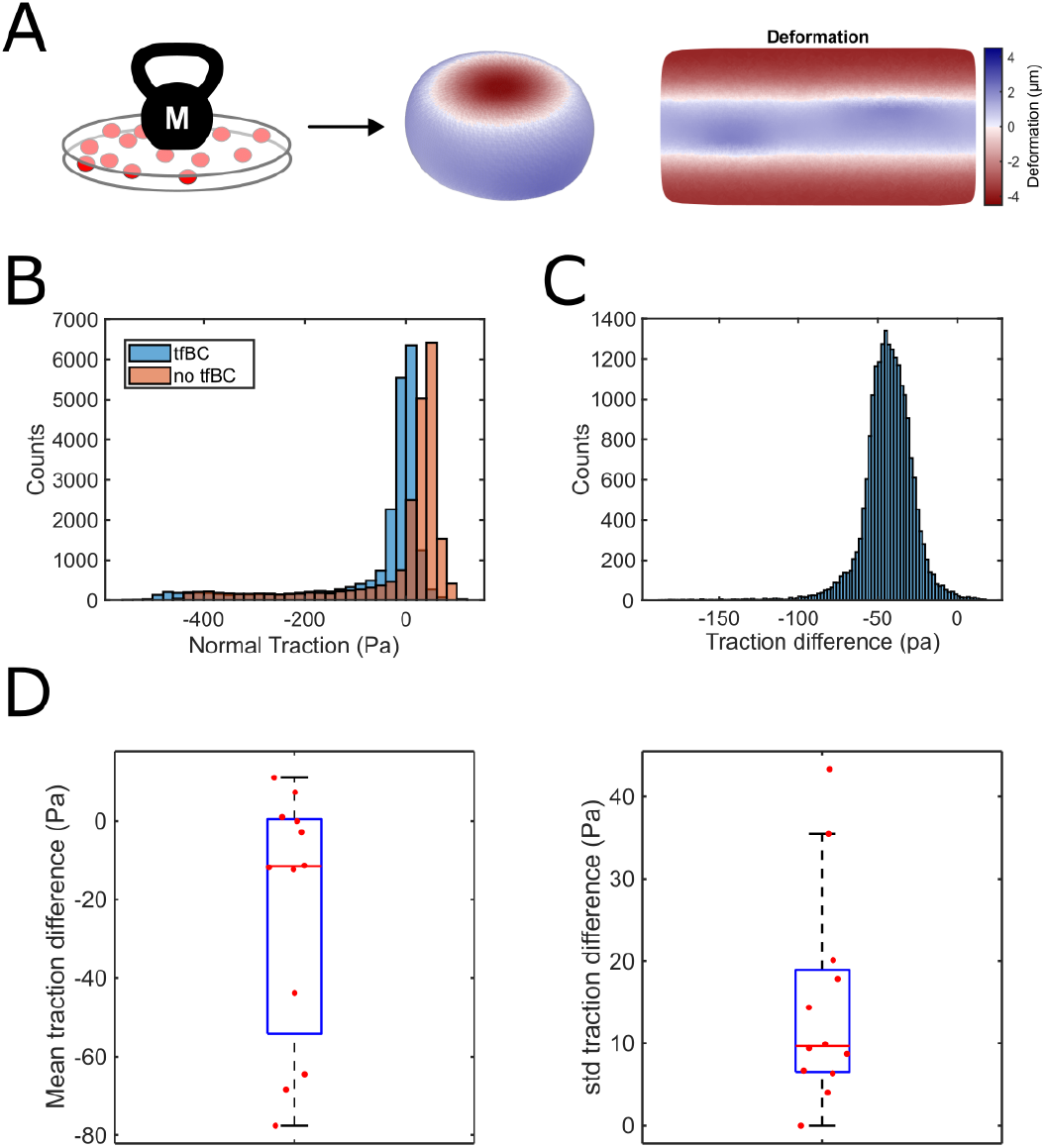
Experiment using a uniaxial load. **A**. Schematic representation of the test experiment. A 10 µL droplet containing microparticles was sandwiched between two glass coverslips. A weight was added on top of the sandwich. Subsequently, the surface of the particles was reconstructed as described showing the predicted flat indentation at the top and bottom where the glass-slides were in contact with the particle, and a particle-expansion in between. **B**. Deformations were subsequently translated into traction as described in the Materials & Methods paragraph. Here we compared the resulting normal traction for two constraints applied: (i) minimization of the total elastic energy (“no tfBC”), and (ii) minimization of the total elastic energy and the traction for which the deformation was positive (“tfBC”). The latter describes the proper situation, as traction occurs only at the coverslip-particle interface and must vanish elsewhere. We noticed that the shapes of the two traction distributions were very similar, with a pronounced peak at low absolute values, and a broad shoulder towards large negative traction (compression). Applying the traction free boundary condition (tfBC, (ii)) resulted in an overall shift of the traction distribution by -41 Pa when compared to the non-tfBC case. With the tfBC applied, the main peak was centered at zero, and was characterized by a width of *σ* = 23 Pa, given by our measurement accuracy. Condition (ii) was used throughout the current study. **C**. The difference between normal traction of the two minimization methods appeared to be near-Gaussian, of mean -43 Pa and width of 18 Pa. **D**. Analysis was performed on 12 microparticles. For all, there was a shift of the traction-distributions, while the shape of the distributions did not change. The shift for the two minimization procedures ranged from 0 to -80 Pa, with a median width of *σ* ≈ 10 Pa. This value is less than our traction resolution as characterized by the width of the low absolute value peak of ≈ 30 Pa, indicating that applying the tfBC mainly generates an offset rather than a shape-change of the distribution.

**Figure S3:**
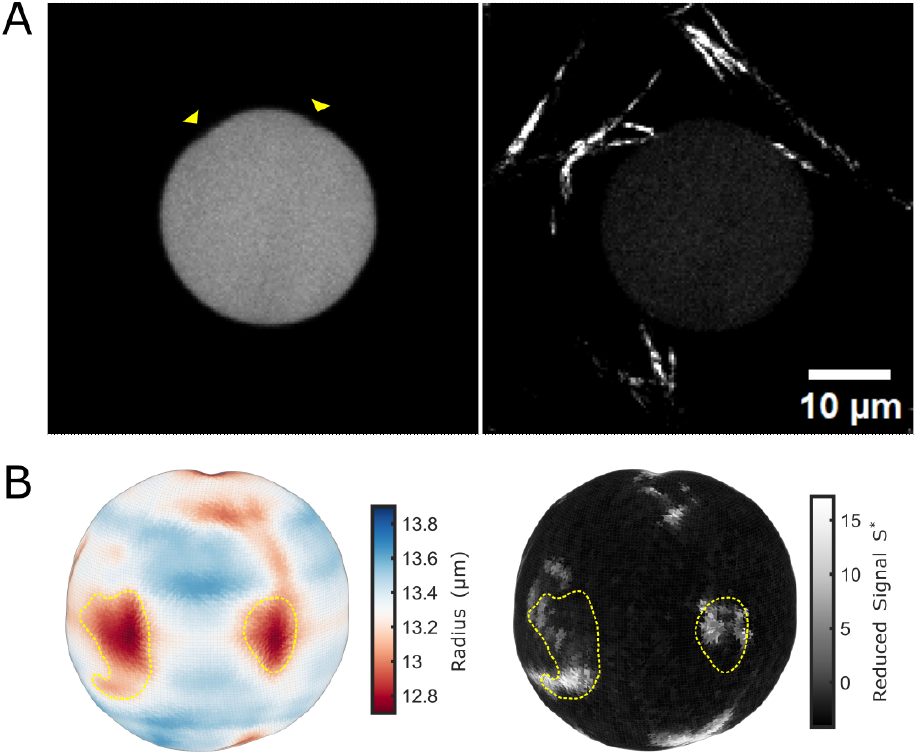
Thick collagen fibers apply local stresses on the microparticles. **A**. TexasRed-Cadaverine labelled microparticle (left) and reflection microscopy of the surrounding collagen fibers (right). Thick collagen fibers significantly indent the particle (yellow arrows). The dense thin-fiber collagen network was not visible at those experimental settings. **B**. Quantification of the microparticle shape (left) and collagen signal on the microparticle surface (right). The indentations (yellow outline) coincide with regions of high collagen signal.

**Figure S4:**
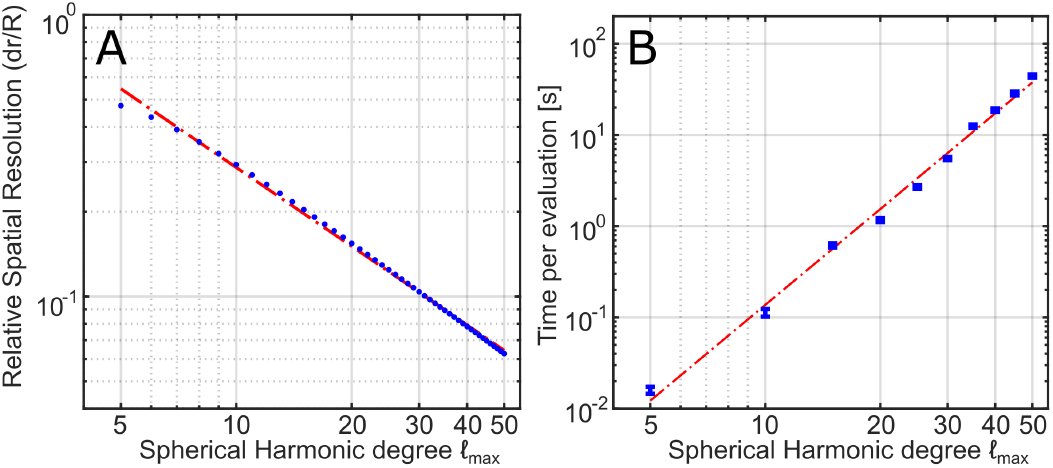
Trade-off between spatial resolution and computational time. With increased harmonics degree *l*, the relative spatial resolution, *dr/R* decreases (**A**) while the time required per computation goes up (**B**). Power law fit is displayed in red. The slope of the power-law fit is, *m* ≈ 3.4.

**Figure S5:**
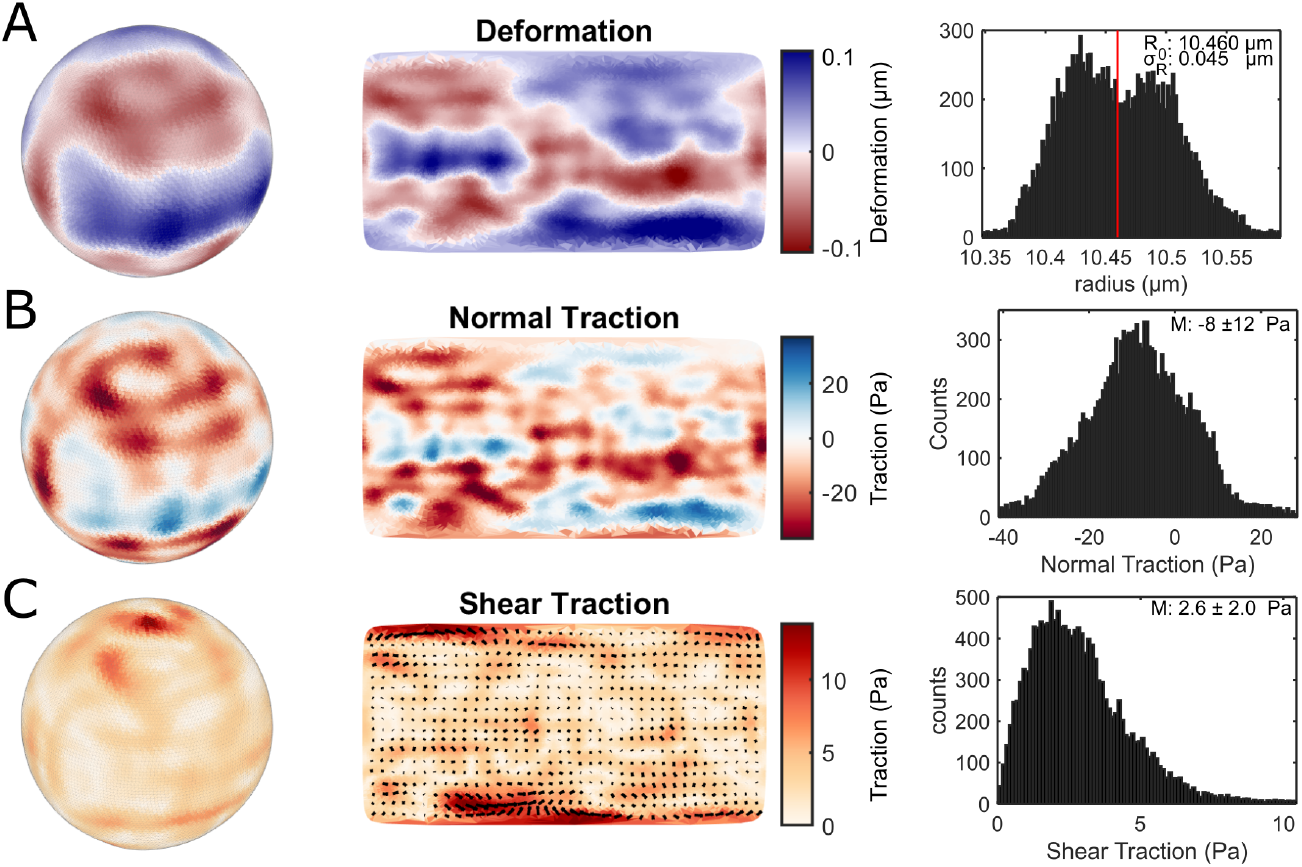
Edge localization accuracy of a particle suspended in a collagen gel. **A**. Deformation field. The standard deviation of the surface deformation is 45 nm rms. **B**. Normal traction field. Its standard deviation is 12 Pa rms. **C**. Shear traction field. Its standard deviation is 2.0 Pa rms.

**Figure S6:**
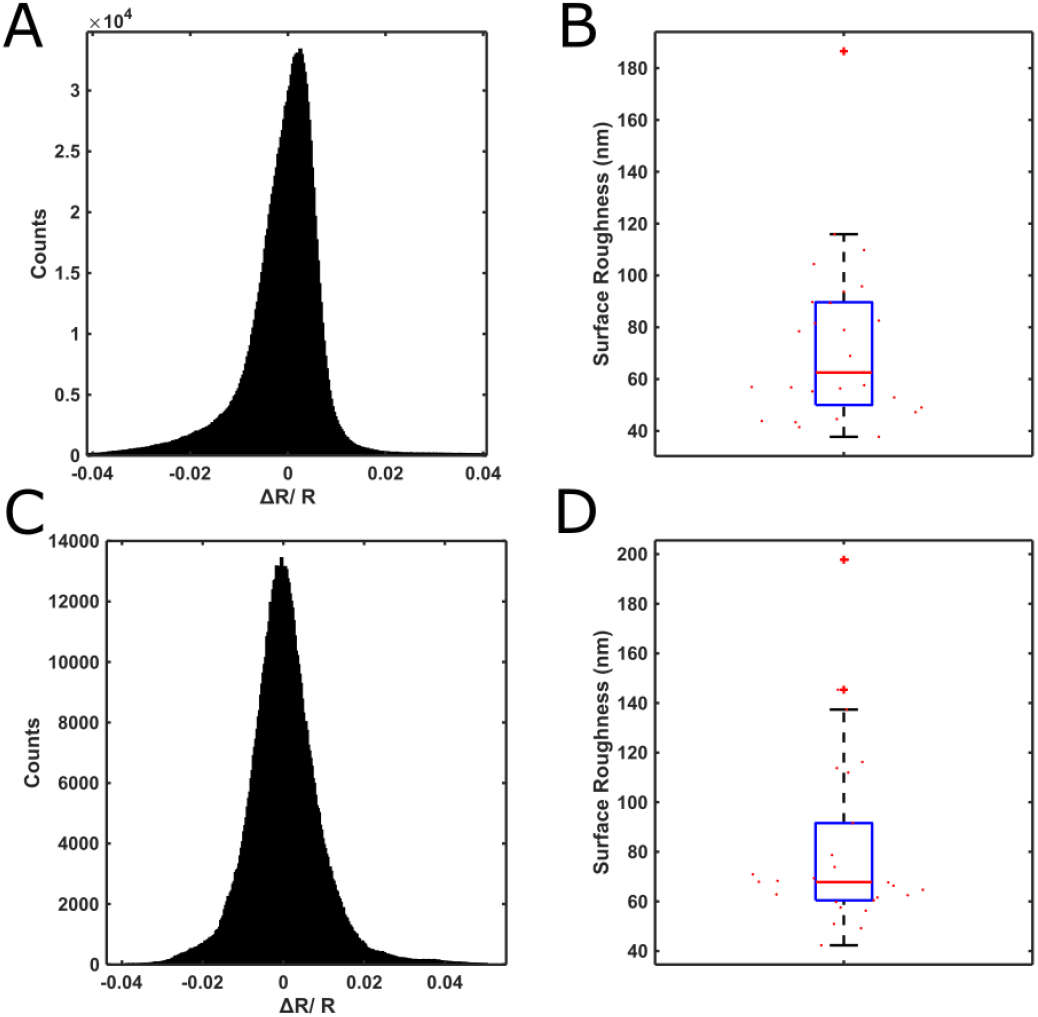
Particles shape reconstruction in ideal (particles in buffer), and realistic (particles in tumoroid-embedded gels far off tumoroids) conditions. **A**. Combined histogram of reduced deformations Δ*R/R* normalized to the radius *R* of 27 microparticles in ideal conditions; on a glass coverslip without cells or collagen. The standard deviation is *σ* = 0.0095 rms. **B**. Boxplot of the individual root mean squared (rms) surface roughness of the distributions from (A). The median rms was 62.5 nm. **C**. Combined histogram of normalized deformations of 26 microparticles in realistic conditions; suspended in a collagen matrix in the absence of cells; *σ* = 0.0085 rms. **D**. Boxplot of the individual rms surface roughness of the distributions from (C). The median rms was 67.8 nm, almost the same as in ideal conditions.

**Figure S7:**
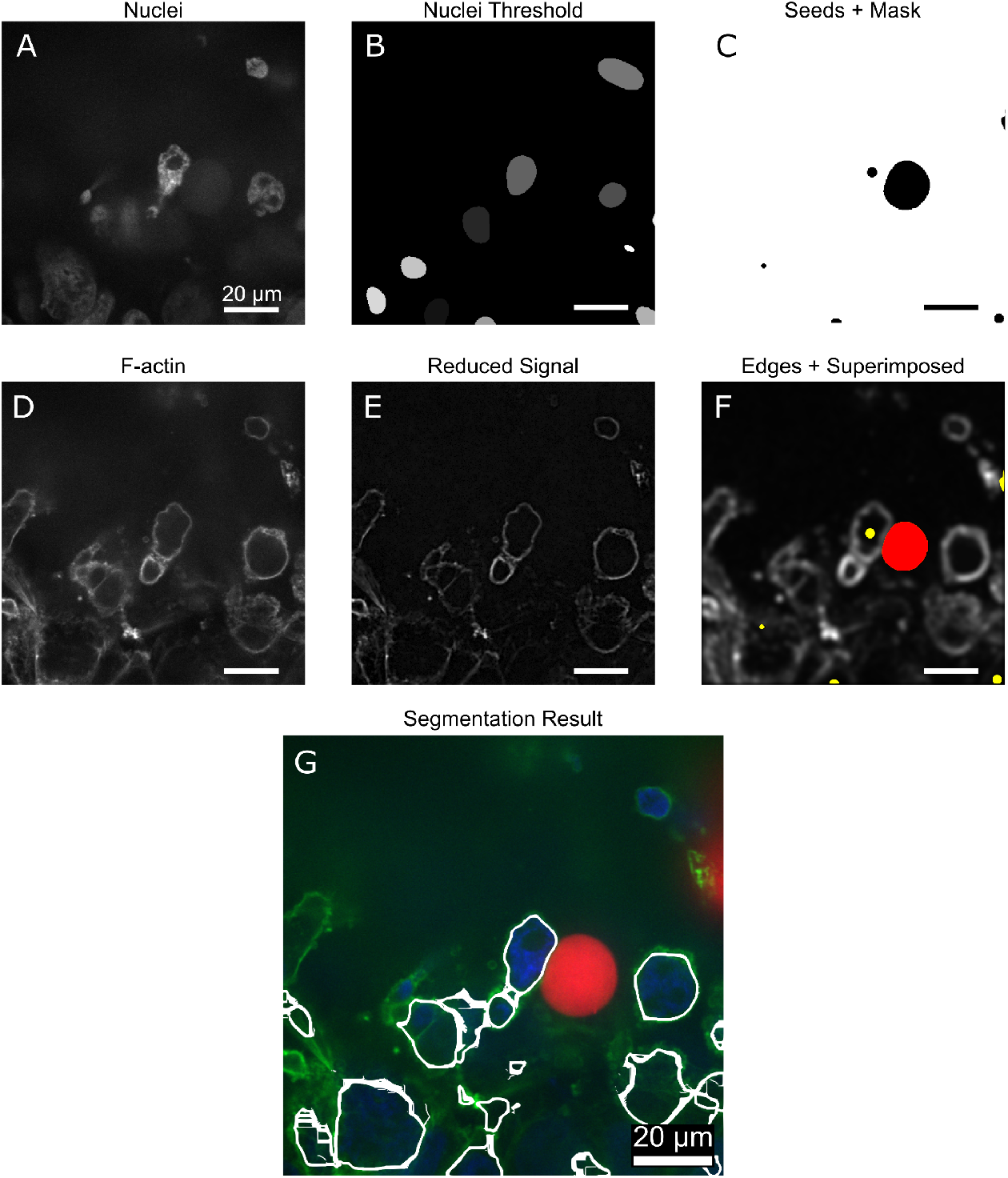
Segmentation method. **A**. Cell were identified based on their nuclei. **B**. A Gaussian filter was applied and nuclei were detected using an adaptive threshold. **C**. The nuclei centers function as seeds for the segmentation. Note that this is a single slice of a 3d image cube. Hence, not all seeds are visible in this slice. Additionally, the particle volume was excluded. **D**. Cell boundaries were identified by labelling of F-actin. **E**. Salt-and-pepper noise was reduced by applying an adaptive median filter. **F**. Using the Sobel operator, edges were found. A Gaussian filter was applied to merge edges originated from the same line segment. Subsequently, the image was superimposed with the seeds (yellow) and the particle (red). **G**. Overlay of the detected cell edges (white) and the composited image. All scale bars are 20 µm.

**Figure S8:**
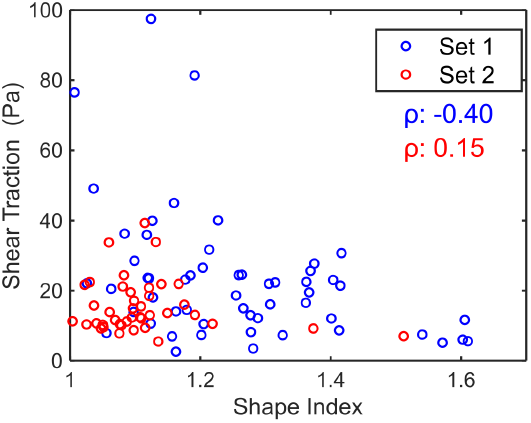
Shear traction appeared to be uncorrelated to the cell shape in two independent experiments. Correlation coefficients are displayed by *ρ*.

